# Effects of sodium butyrate and rosemary leaves on performance, biochemical parameters, immune status, and carcass traits of broiler chickens

**DOI:** 10.1101/2024.02.14.580269

**Authors:** Hamed Yahya Saifan, Mostafa Abbas Shalaby, Khaled Abo-EL-Sooud, M.A. Tony, Aya M. Yassin

## Abstract

Effects of sodium butyrate (SB) and rosemary leaves (RL) on growth performance, biochemical profile, immune status, and carcass traits of broiler were evaluated. Fifty-one-day old Hubbard chicks (unsexed) were purchased from Al-Ahram Company for Poultry, Egypt and reared on floor pens in a private farm. The chicks were weighed on arrival and assigned at random into five equal groups, with four replicates each (25 chicks/replicate). Group 1 was fed on a broiler diet without any additions and used as a control. The diets of groups 2 and 3 were supplemented with 500 g/ton SB and 4 kg/ton RL, respectively. In group 4, the diet was enriched with 250g/ton SB plus 2 kg/ton RL. Chicks in group 5 were fed on a diet fortified with 500 g/ton SB plus 4 kg/ton RL. Body weight gain (BWG) and feed efficiency ratio (FER) were determined weekly. Biochemical indexes, immune system function, and carcass traits were determined. The results revealed that supplementing broiler diet with 500g/ton SB plus 4 kg/ton RL increased BWG and FER of birds. This diet decreased serum levels of aspartate aminotransferase, (AST) alanine aminotransferase (ALT), total cholesterol (TC), triglycerides (TG), malondialdehyde (MDA and increased superoxide dismutase (SOD), catalase (CAT), and immunoglobulins. It also boosted immunity as it elevated phagocytic activity, phagocytic index, lysozyme activity, and nitric oxide (NO) concentrations. Antibody titers against Newcastle disease virus were elevated. It can be concluded that supplementing broiler diet with 500g/ton SB plus 4 kg/ton RL improve performance, normalize biochemical profile, and boost immunity.

## INTRODUCTION

Egypt is suffering from a severe shortage of poultry meat, which is a popular food for most Egyptians. Chicken white meat is a favorite for most Egyptians due to its affordable price when compared to red beef meat. It is estimated that Egypt’s chicken meat production will be approximately 1.59 million tons in year 2023 **(Abdelli *et al*. 2021)**. The cost of feed ingredients such as cereal grains (maize, sorghum, and barley) tends to increase because of fluctuations of prices and difficulties in the importation process. Antibiotics have been used as a traditional additive for promoting growth performance and health of poultry. The risk of bacterial resistance development and the incidence of drug residues that pose great risks to human health leading to antibiotic withdrawal from use in poultry nutrition **(Ricke *et al*. 2020)**.

Feed additives are utilized in chicken nutrition for a number of reasons, such as augmenting the safety and quality of the feed, improving the quality of animal-derived byproducts, and enhancing the growth and carcass characteristics. Antimicrobial medications **(Alagawany *et al*. 2021)**; acidifiers **(Ricke *et al*. 2020)**; antioxidants **(Hashemi and Davoodi, 2011)**; antimycotoxin **(Olivera *et al***. 2015); prebiotics, probiotics, and phytogenic additives **(Abd El-Hack *et al*. 2017 and Al-Khalaifah, 2018)** are the most often used feed additives for poultry diets.

For the production of chicken, sodium butyrate (SB), the sodium salt of butyric acid, is a short chain fatty acid that is frequently utilized. According to a prior study **(Lan *et al*. 2020)**, it was reported that feeding broilers a basal diet supplemented with SB enhanced growth performance, liver function, antioxidant capacity, carcass features, and meat quality. Particularly for birds receiving treatments for coccidiosis, SB may be a helpful tactic to support gut health **(Sadurní *et al*. 2022)**.

The growth performance of broilers related to the regulation of intestinal flora was enhanced by the addition of 1000 mg/kg of protected sodium butyrate (CSB) to their diet **(Zhao *et al*. 2022)**. A considerable improvement in body weight gain, feed intake, feed conversion ratio, were observed when SB and the antibiotic bacitracin methylene disalicylate (BMD) were combined **(Kumar *et al*. 2023)**.The effects of xylo-oligosaccharide (XOS) and chemically protected sodium butyrate (CSB), alone and in combination, enhanced broiler growth performance and produced anti-inflammatory and antioxidant properties **(Deng *et al*. 2023)**.

Rosemary (*Rosmarinus officinalis*) is known to contain high levels of saponin, tannin, and resin. It was recognized that rosemary herb produced antibacterial and antioxidant properties **(Mathlouthi *et al*. 2012 and Moreno *et al*. 2018)**. Immune system, meat quality, and production performance of broiler chickens have all been shown to benefit from the addition of rosemary to their diet **(Ghazalah and Ali, 2008)**. Additionally, **Yesilbag *et al*. (2011)** documented the benefits of using rosemary oil to laying quails. On the other hand, **Loetscher *et al*. (2013)** reported that the addition of rosemary to chicken diets do not had a significant impact on production and slaughter traits. However, the prevailing opinion is that rosemary has a significant potential in the nutrition of poultry, but it is important to choose the proper source, form, concentration, and mode of practical application in broiler diets.

## MATERIAL AND METHODS

### Ethical Approval

The current study was approved according to the Institutional Animal Care and Use Committee (IACUC), Faculty of Veterinary Medicine, Cairo, University, Egypt, with reference number: Vet CU 09092023791, dated at 9/ 9/ 2023.

### Feed Additives

Sodium butyrate (SB), trade name CM3000®, is a commercial 30% spherical granule-coated feed supplement. It is the sodium salt of butyric acid with a chemical formula of C4H7NaO2. It is a short-chain fatty acid that is released slowly and continuously in both the small and large intestine of poultry. CM3000® is manufactured by Hangzhou King Techina Feed Co., Ltd., China. It was added to the broiler basal diet at a concentration of 500 g/ton feed as reported by **Sikandar *et al*. (2017)**.

Rosemary, *Rosmarinus* officinalis L. powder (RL), Family *Lamiaceae*, is used as a feed additive for poultry based on a natural source. Rosemary is a well-known traditional medicine and cooking spice. It acts as a useful poultry dietary supplement for boosting immunity. Essential oils of rosemary represents an effective growth promoter in broilers. Rosemary leaves were purchased from a local market (Haraz for Herbs, Medicinal Plants, Spices, and Natural Oils) Cairo, Egypt. The dry leaves were finely powdered using an electrical mill and then added to the basal diet at a concentration of 4 kg/ton according to **Al-Kassie *et al*. (2008)**.

### Experimental Chicks and Diets

Fifty one-day-old Hubbard chicks (unsexed) were purchased from Al-Ahram Company for Poultry, Giza, Egypt. After weighting on arrival, the chicks were randomly assigned into five equal groups with four replicates for each (25 birds/replicate). Group 1 was fed on broiler basal diet without any additions and served as a control. Chicks in groups 2 and 3 were fed on basal diet supplemented with 500 g/ton of SB and 4 kg/ton of RL, respectively. The group 4 was fed on basal diet enriched with 250 SB and 2 kg/ton of RL. Chicks in the group 5 were fed on basal diet fortified with 500 g/ton SB and 4 kg/ton RL. On day 35 of broiler age, the growth performance, biochemical profile, immune status, and carcass traits were evaluated. The experiment was conducted on floor pens at a private farm at Giza, Egypt. Vaccination program for all experimental groups of birds included protection against Newcastle disease (ND), infectious bursal disease (IBD, Gumboro) and infectious bronchitis (IB). To meet the nutrient requirements of Hubbard broilers, diets comprising of corn-soybean meal and basal components were prepared according to Hubbard manual catalogue (2018). During the experimental period (35 days) supplemented diets in the form of mash-type for three stages: starter, grower, and finisher were provided and water was offered *ad labium*.

### Growth Performance

On day 35, the chicks were weighed and their daily feed intake (FI) was reported throughout the experiment period. Body weight gain (BWG) was computed weekly. The feed conversion ratio (FCR) was calculated (g) as mentioned by **Kidane *et al*. (2017)**.

### Collection of Blood

On the 35th day, thirty chickens from each group were chosen at random and 5ml of blood was withdrown from the brachial wing vein into dry plain tubes. Blood was allowed to clot at room temperature. To obtain clear serum, the clots were removed by centrifugation at 2,000–3,000 X g for 15 minutes in a refrigerated centrifuge. The serum samples were poured in Eppendorf tubes and kept in a refrigerator until biochemical analysis.

### Biochemical Analysis

Serum samples were collected to determine aspartate aminotransferase (AST) and alanine aminotransferase (ALT) activities according to the method of **Bergmeyer *et al*. (1978**). Total protein (TP) was determined using the biuret method according to **Zheng *et al*. (2017)**. Total protein (g/L) was calculated using commercially accessible diagnostic kits and an automated biochemical analyser (Alizé). Serum uric acid (UA) and creatinine levels were estimated as described by **Lorentz and Brendt, (1967) and Agbafor *et al*. (2015)**, respectively. Serum total cholesterol (TC) was determined calorimetrically according to **Allain *et al*. (1974)** and triglycerides (TG) according to **Wahlefeld, (1974)**. The level of serum malondialdehyde (MDA) was measured according to the method described by Ohkawa et al., (1979). The activities of serum superoxide dismutase (SOD) and catalase (CAT) enzymes were respectively assessed according to **Nishikimi *et al*. (1972) and Aebi, (1984)** using a spectrophotometer. The zone electrophoresis method was used to separate the serum protein fractions on an agarose gel plate according to **Tothova *et al*. (2019)**.

### Immune Status

The phagocytosis test was accomplished following the procedure of Bos and de Souza, (2000). Phagocytic activity (PA) is the proportion of phagocytic cells that have been engulfed by Candida albicans yeast cells, expressed as a percentage. Phagocytic index (PI) is the number of yeast cells phagocytized divided by the number of macrophage phagocytic cells. Serum samples were withdrawn from brachial wing vein at day 7 and day 21 post-vaccination with Newcastle diasese virus. The lytic activity of lysozyme against the cell wall of Micrococcus lysodeikticus was used as a substrate in the lysozyme assay (**LA**) method. To conduct this assay, an agarose gel plate lysate method was employed, following the protocol outlined by **Peeters and Vantrappen, (1977)**. The lysozyme concentration was determined by generating a logarithmic curve with a standard lysozyme solution. nitric oxide (NO) assay was accomplished in accordance with **Sun *et al*. (2003)** using Griess reaction assay after removing protein via mixture of ZnSO4 and NaOH. The absorbance at 540 nm exhibits a linear correlation with the concentration of NO present in the sample. Antibody titres against newcastle disease virus using Hemagglutination inhibition (HI) test according to **Beard, (1989)** were measured. On day 7 and day 21, six broilers from each group were randomly selected for estimation of HI antibody titres. Briefly, blood samples (2 ml from each broilers) were withdrawn from the brachial vein into non-heparinized vacuum tubes (Becton Dickinson Vacationer Systems, Franklin Lakes) and allowed to clot at 4°C for 2 hours. The serum was separated by centrifugation at 3,000 g for 15 min, and stored at −20°C for HI anti-body assay. After the serum was inactivated at 56°C for 30 min, twofold serial dilution were made in a 96-well V-shaped bottom microtitre plate containing 50 μl of calcium and magnesium-free (CME) phosphate buffered saline (PBS), in each well then 50 μl of NDV antigen (4 HA units) was added into all wells except the last row kept as controls. Serum dilutions ranged from 1:21 to 1:212. The plate was incubated at 37°C for 10 min, then 50 μl of 1% erythrocytes suspension was added to each well and incubated for 30 min. A positive serum, a negative serum, erythrocytes and antigens were also included as controls. The last wells which caused complete inhibition was considered as the endpoint. The geometric mean titres was expressed as reciprocal log 2 values of the last dilution and the absence of hemagglutination is a positive result. The enzyme linked immunosorbent assay (ELISA) technique, as outlined by **Engvall and Perlmann, (1971)** was used to determine levels of serum IgG and IgM.

### Carcass Charactristics

At the end of experimental period, thirty birds were randomly chosen from each group and prepared for slaughtering. The birds were fasted for 12 hours then slaughtered by bleeding of the jugular vein. Once slaughtered, the birds were defeathered and eviscerated. Heart, liver, spleen, thymus, bursa, and abdominal fat were removed and weighed on a digital scale. Head and offals were removed and the remaining carcasses were weighed to obtain the ready-to-cook carcass weight. Using this weight, the carcass dressing yield percentage (dressing %) was then calculated according to **Rosa *et al*. (2007)**.

### Statistical Analysis

Data were recorded as means ± SD. To analyse the data, IBM SPSS® version 19 software was utilized on a personal computer (2010). The means ± SD were compared with a one-way ANOVA test, with a significance level of P<0.05, and the Post Hoc Duncan test was then applied according to **Snedecor and Cochran, (1986)**.

## RESULTS

### Growth Performance

The present results revealed that supplementation of a basal diet with sodium butyrate (SB) and rosemary leaves (RL) alone and in combination increased body weight gain (BWG) and feed conversion ratio (FCR) on day 35 of age of broilers as recorded in Table 1.

**Table 1.** Effect of sodium butyrate and rosemary leaves on the growth performance of broiler chickens (n=30 birds)

| Groups<br>Parameters | G1 | G2 | G3 | G4 | G5 |
| --- | --- | --- | --- | --- | --- |
| Initial body weight<br>on day 28 (g) | 1355.16<br>±0.12<sup>a</sup> | 1390.11<br>±0.12<sup>c</sup> | 1410.11<br>±0.12<sup>c</sup> | 1422.30<br>±0.18<sup>b</sup> | 1443.91<br>±0.15<sup>b</sup> |
| Final Body weight<br>on day 35 (g) | 1885.42<br>±0.68<sup>a</sup> | 1986.58<br>±0.53<sup>c</sup> | 2050.37<br>±0.55<sup>c</sup> | 2090.80<br>±0.61<sup>b</sup> | 2120.17<br>±0.62<sup>b</sup> |
| Weight gain (g) | 530.26<br>±0.15<sup>a</sup> | 596.74<br>±0.18<sup>c</sup> | 640.26<br>±0.15<sup>c</sup> | 668.50<br>±0.22<sup>c</sup> | 676.26<br>±0.21<sup>b</sup> |
| Feed intake (FI) (g) | 876.90<br>±0.45<sup>a</sup> | 998.50<br>±0.60<sup>d</sup> | 1100.31<br>±0.60<sup>d</sup> | 1160.69<br>±0.55<sup>c</sup> | 1185.77<br>±0.58<sup>b</sup> |
| Feed conversion<br>ratio (FCR) | 1.65 | 1.67 | 1.71 | 1.73 | 1.75 |

Means ± SD with different letter superscripts are significant at $P \leq 0.05$
G1= Group of chicks fed on basal diet (BD)
G2= Group of chicks fed on BD supplemented with 500 g/ton of sodium butyrate (SB)
G3= Group of chicks fed on Group fed on BD supplemented with 4 Kg/ton of rosemary leaves (RL)
G4 = Group of chicks fed on BD supplemented with 250 g/ton of SB + 2 kg/ton of RL
G5 = Group of chicks fed on BD supplemented with 500 g/ton of SB + 4 kg/ton of R

### Biochemical Profile

Table 2 shows that fortification of a basal diet with SB and RL alone and in combination significantly decreased AST, ALT, TC and TG in serum of broiler chickens.

**Table 2.**
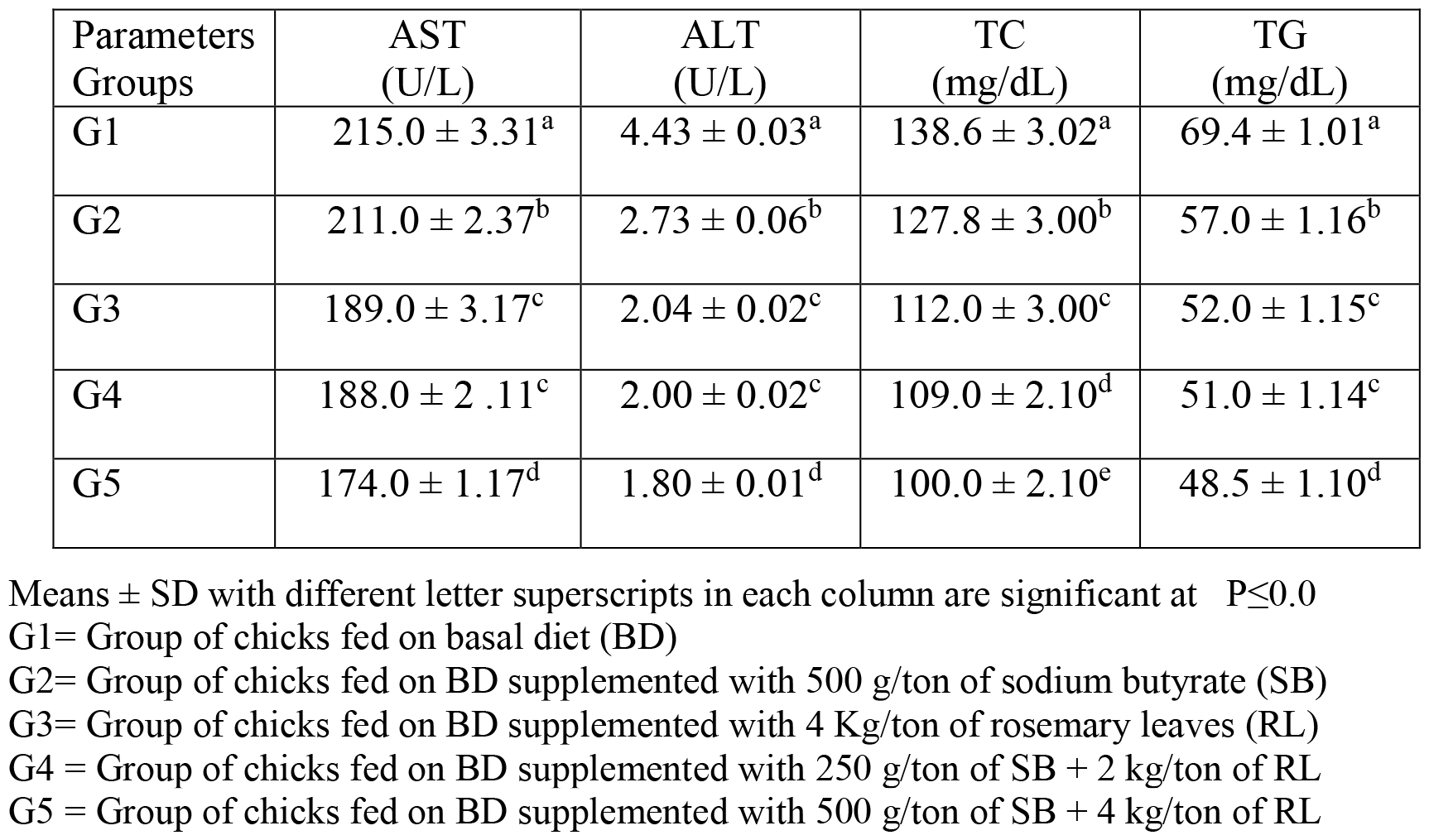
Impact of sodium butyrate (SB) and rosemary leaves (RL) alone and in combination on serum concentration of liver enzymes (AST and ALT), total cholesterol (TC), and triglycerides (TG) on day 35 of age of broiler chickens (n=30 birds).

| Parameters<br>Groups | AST<br>(U/L) | ALT<br>(U/L) | TC<br>(mg/dL) | TG<br>(mg/dL) |
| --- | --- | --- | --- | --- |
| G1 | 215.0 ± 3.31<sup>a</sup> | 4.43 ± 0.03<sup>a</sup> | 138.6 ± 3.02<sup>a</sup> | 69.4 ± 1.01<sup>a</sup> |
| G2 | 211.0 ± 2.37<sup>b</sup> | 2.73 ± 0.06<sup>b</sup> | 127.8 ± 3.00<sup>b</sup> | 57.0 ± 1.16<sup>b</sup> |
| G3 | 189.0 ± 3.17<sup>c</sup> | 2.04 ± 0.02<sup>c</sup> | 112.0 ± 3.00<sup>c</sup> | 52.0 ± 1.15<sup>c</sup> |
| G4 | 188.0 ± 2.11<sup>c</sup> | 2.00 ± 0.02<sup>c</sup> | 109.0 ± 2.10<sup>d</sup> | 51.0 ± 1.14<sup>c</sup> |
| G5 | 174.0 ± 1.17<sup>d</sup> | 1.80 ± 0.01<sup>d</sup> | 100.0 ± 2.10<sup>e</sup> | 48.5 ± 1.10<sup>d</sup> |

Means ± SD with different letter superscripts in each column are significant at $P \leq 0.05$
G1= Group of chicks fed on basal diet (BD)
G2= Group of chicks fed on BD supplemented with 500 g/ton of sodium butyrate (SB)
G3= Group of chicks fed on BD supplemented with 4 Kg/ton of rosemary leaves (RL)
G4 = Group of chicks fed on BD supplemented with 250 g/ton of SB + 2 kg/ton of RL
G5 = Group of chicks fed on BD supplemented with 500 g/ton of SB + 4 kg/ton of RL

The present results indicated that addition of SB and RL alone and in combination to the broilers’ basal diet caused significant increases in serum levels of SOD and CAT antioxidant enzymes and a decrease in MDA serum level as recorded in Table 3.

**Table 3.**
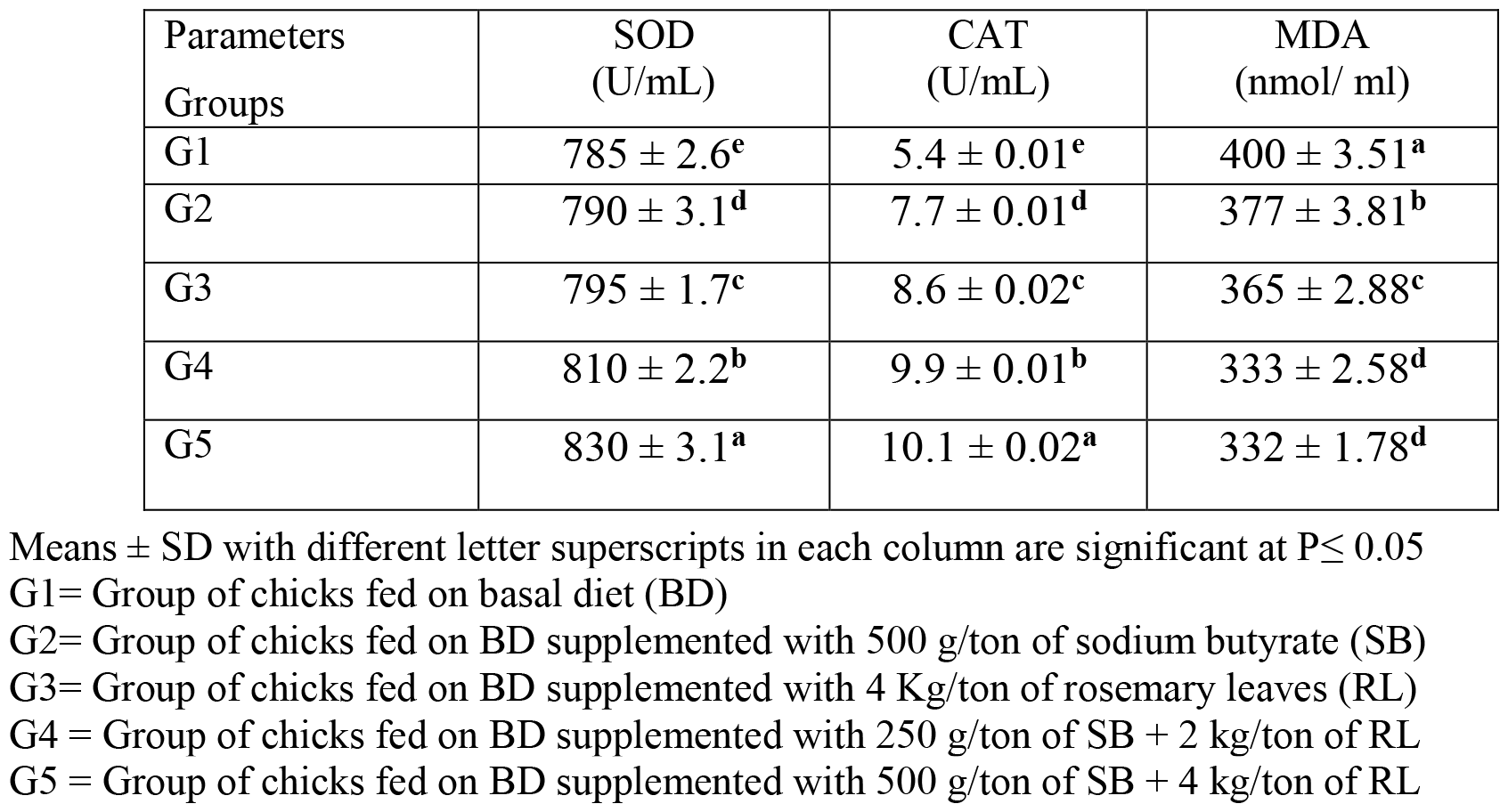
Impact of sodium butyrate (SB) and rosemary leaves (RL) alone and in combination on serum superoxide dismutase (SOD), catalase (CAT), and malondialdehyde (MDA) levels on day 35 of age of broiler chickens (n=30 birds).

The results of this study indicated that supplementation of basal diet with sodium butyrate (SB) and rosemary leaves (RL) alone and in combination significantly increased total proteins (TP), albumin **(Alb)**, globulin (Glb) and albumin/globulin ratio on day 35 of age of broilers as shown in Table 4.

**Table 4.** Effect of sodium butyrate (SB) and rosemary leaves (RL) alone and in combination on serum concentrations of total proteins (TP), albumin (Alb), globulin (Glb) and albumin/ globulin ratio in serum of broilers on day 35 of age of broiler chickens (n=30 birds).

| Parameters Groups | TP (g/mL) | Alb (g/mL) | Glb (g/mL) | Alb/Glb Ratio |
| --- | --- | --- | --- | --- |
| G1 | 32.2 ± 1.08<sup>d</sup> | 13.0 ± 0.9<sup>d</sup> | 20.2± 0.9<sup>d</sup> | 0.643 |
| G2 | 34.1 ± 0.53<sup>c</sup> | 15.9 ± 0.3<sup>c</sup> | 23.1± 0.7<sup>c</sup> | 0.688 |
| G3 | 38.8 ± 0.23<sup>b</sup> | 19.0 ± 0.4<sup>b</sup> | 27.2± 0.6<sup>b</sup> | 0.698 |
| G4 | 39.4 ± 0.23<sup>b</sup> | 20.5 ± 0.2<sup>b</sup> | 28.2± 0.5<sup>b</sup> | 0.726 |
| G5 | 41.1 ± 1.06<sup>a</sup> | 24.4 ± 0.3<sup>a</sup> | 32.5± 0.3<sup>a</sup> | 0.750 |

Means ± SD with different letter superscripts in each column are significant at P≤ 0.05

As elucidated in Fig.1, the addition of sodium butyrate (SB) and rosemary leaves (RL) alone and in combination to a basal diet elevated serum immunoglobulin IgG and IgM concentrations in broiler chickens on day 35 of age.

**Fig. 1.**
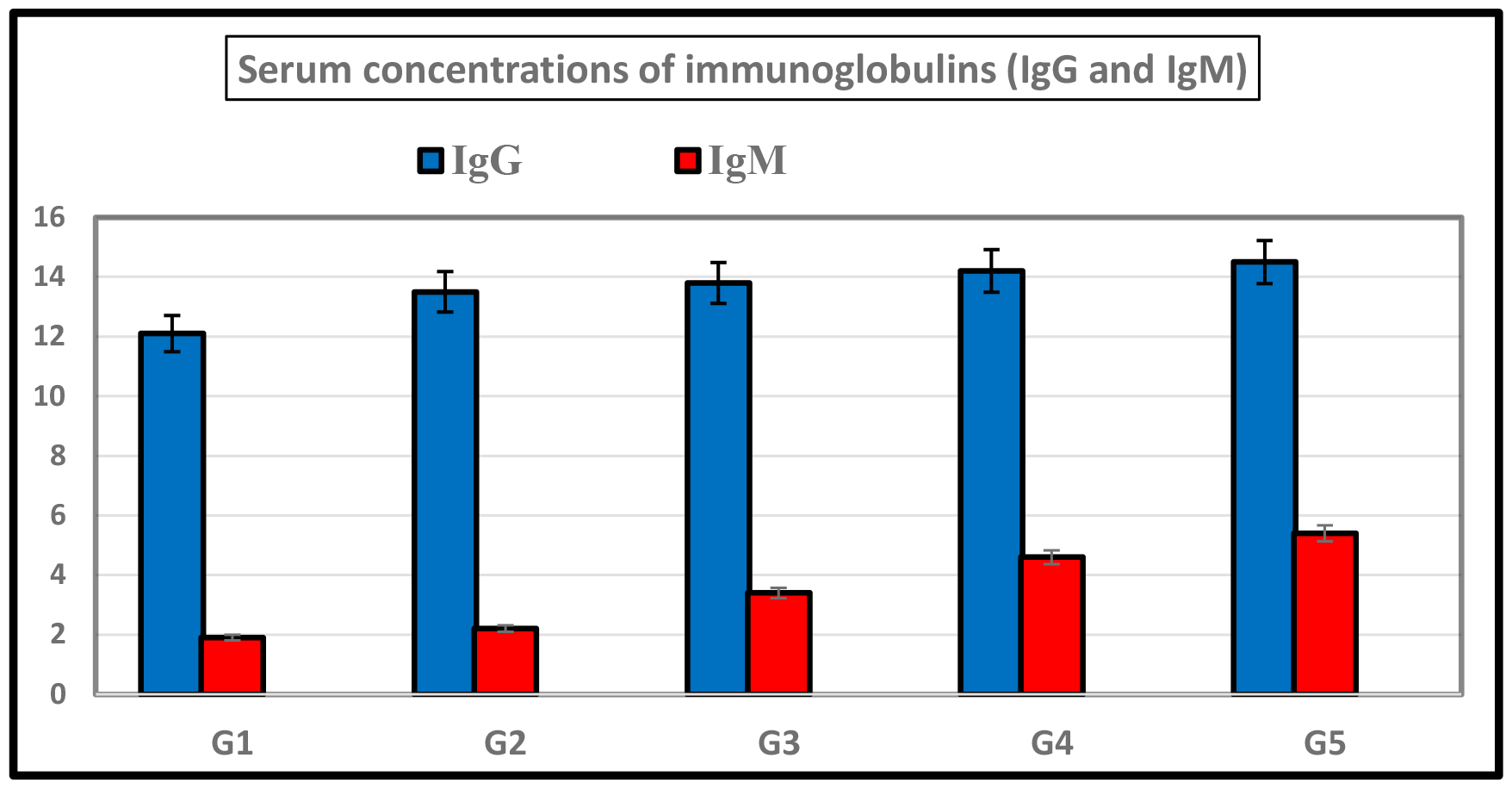
Showing the effect of sodium butyrate (SB) and rosemary leaves (RL), alone and in combination, on serum concentration of IgG and IgM on day 35 of age of broiler chickens (n=30 birds). G1= Group of chicks fed on basal diet (BD) G2= Group of chicks fed on BD supplemented with 500 g/ton of sodium butyrate (SB) G3= Group of chicks fed on BD supplemented with 4 Kg/ton of rosemary leaves (RL) G4 = Group of chicks fed on BD supplemented with 250 g/ton of SB + 2 kg/ton of RL G5 = Group of chicks fed on BD supplemented with 500 g/ton of SB + 4 kg/ton of RL

As recorded in Table 5, the addition of sodium butyrate (SB) and rosemary leaves (RL), alone and in combination, increased macrophage phagocytic activity (PA), phagocytic index (PI), lysozyme activity (LA) and serum nitric oxide (NO) concentration in broiler chickens.

**Table 5.**
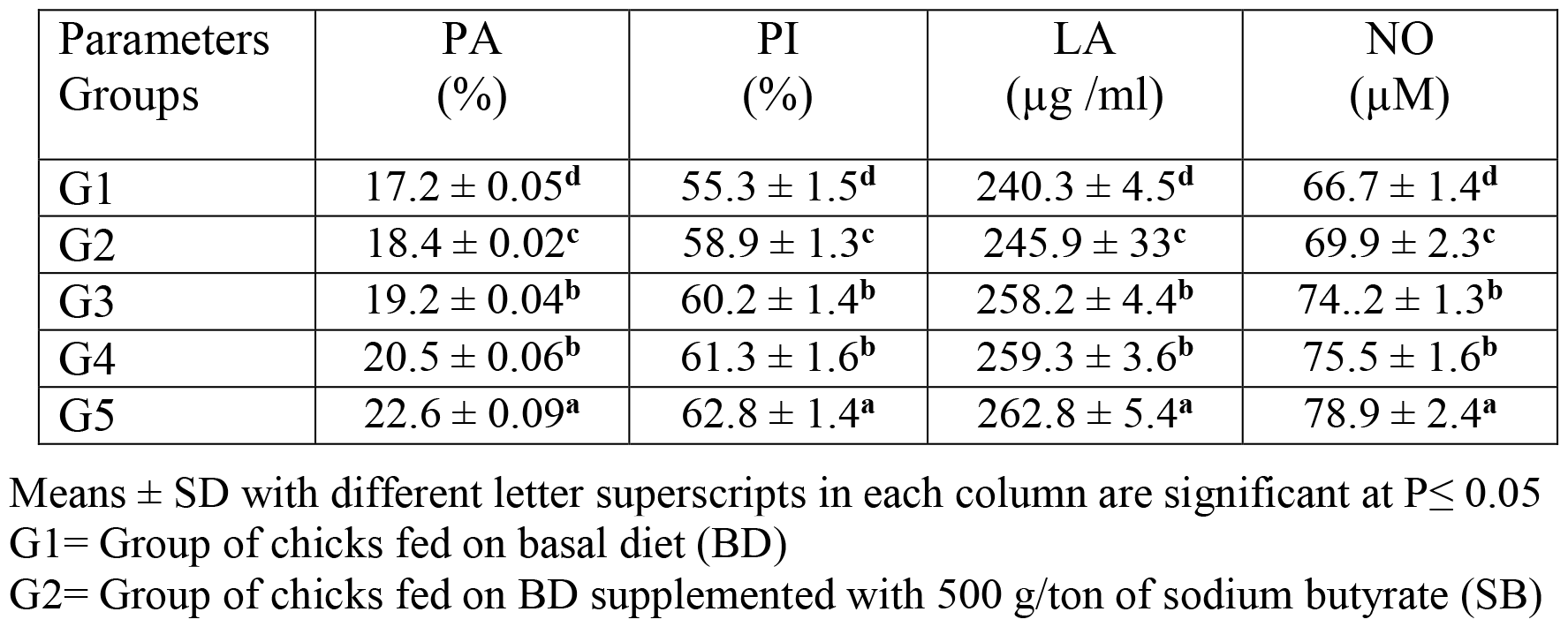
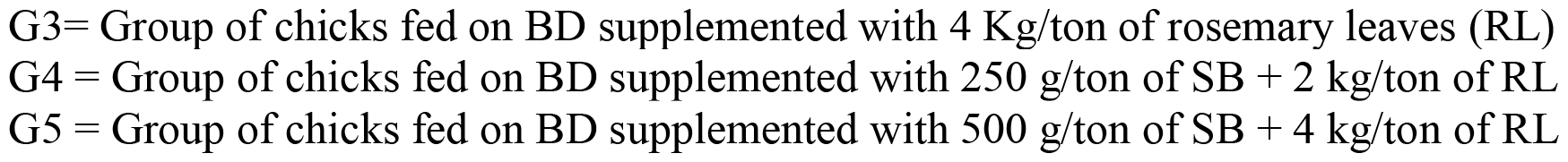
Effect of sodium butyrate (SB) and rosemary leaves (RL) alone or in a mixture on phagocytic activity (PA), phagocytic index (PI), lysozyme activity (LA) and serum nitric oxide (NO) concentration on day 35 of age of broiler chickens (n=30 birds).

Results of the current study revealed that hemagglutination inhibition (HI) antibody titers against NDV were significantly increased in sera collected on day 7 post vaccination with Hitchner B1 strains and on day 21 post vaccination with Lasota strains. It was found that Lasota vaccine had a good potency against newcastle disease in broilers as demonstrated in Fig.2.

**Fig. 2.**
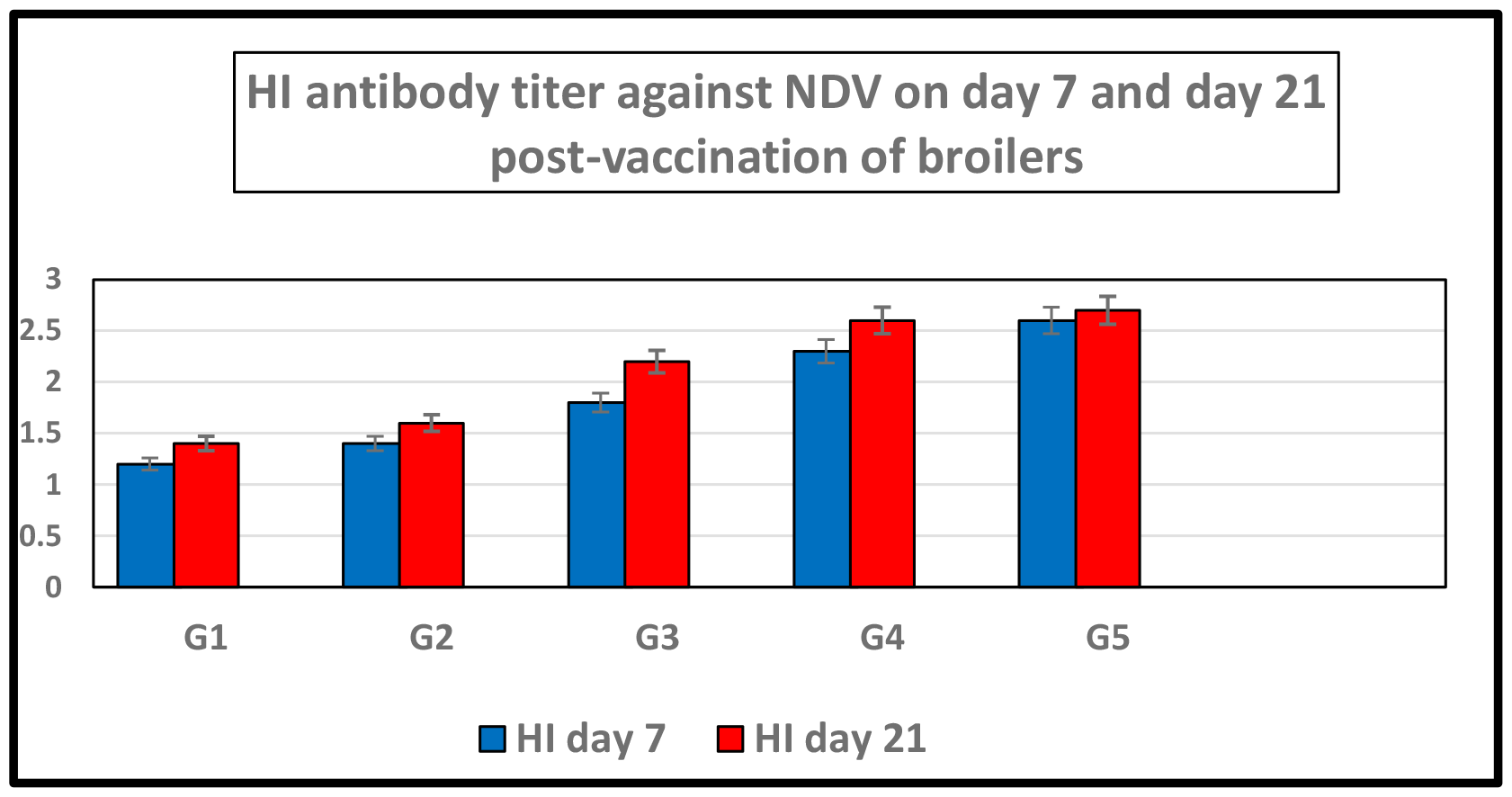
Showing the effect of sodium butyrate (SB) and rosemary leaves (RL), alone and in combination, on hemmaglutination inhibition antibody titer against NDV on day 7 and day 21 post-vaccination of broilers (n=30 birds). G1= Group of chicks fed on basal diet (BD) G2= Group of chicks fed on BD supplemented with 500 g/ton of sodium butyrate (SB) G3= Group of chicks fed on BD supplemented with 4 Kg/ton of rosemary leaves (RL) G4 = Group of chicks fed on BD supplemented with 250 g/ton of SB + 2 kg/ton of RL G5 = Group of chicks fed on BD supplemented with 500 g/ton of SB + 4 kg/ton of RL

As depicted in Table 6, the addition of sodium butyrate (SB) and rosemary leaves (RL), alone and in combination, increased live and carcass weights. The dressing percent (DP %) ranged from 71% to 72.77%. There were significant increases in weights of spleen, thymus and bursa and a decrease of abdominal fat. Non-significant changes were reported in weights of heart and liver.

**Table 6.**
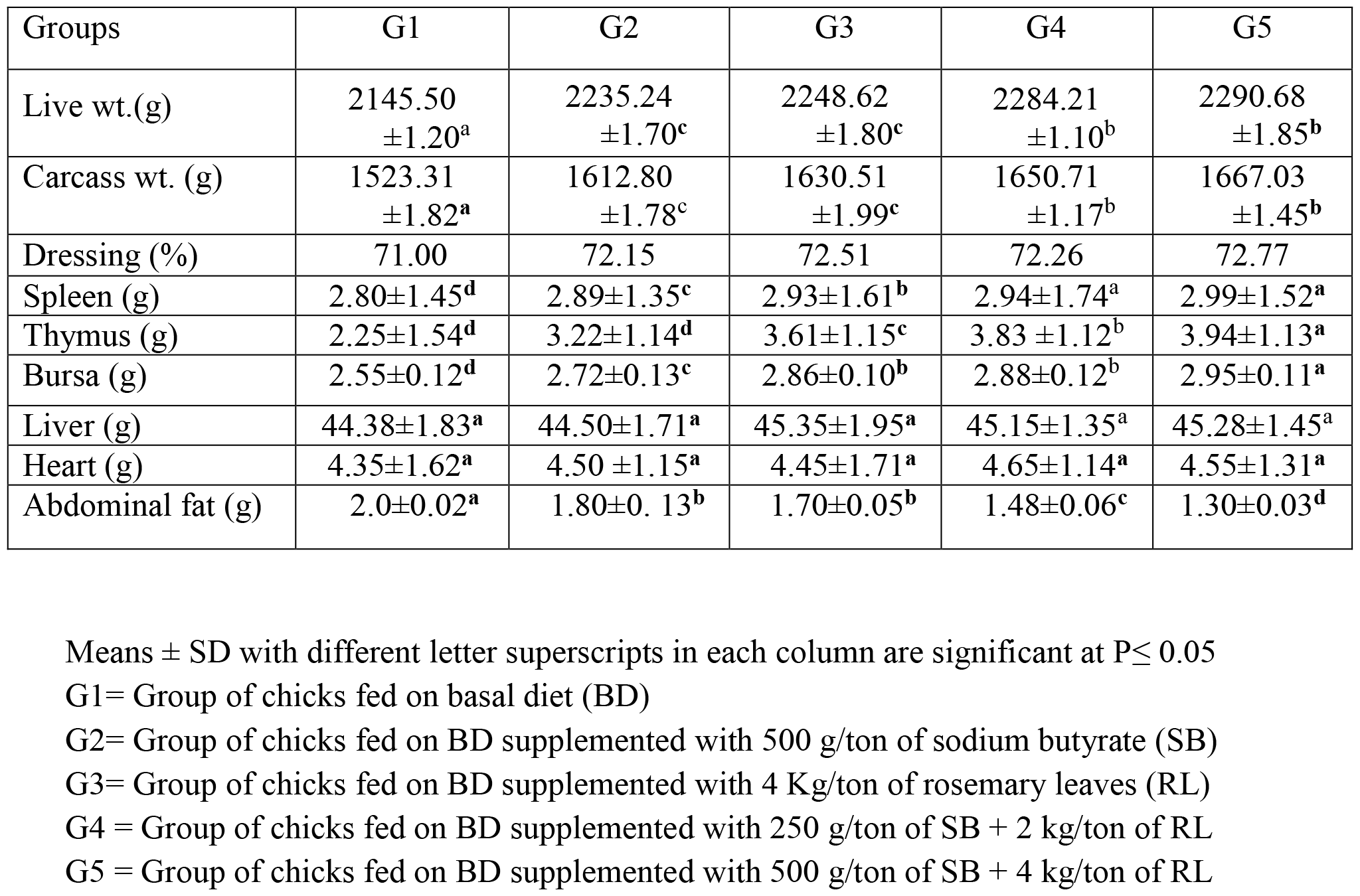
Effect of sodium butyrate (SB) and rosemary leaves (RL) alone and in combination on carcass traits on 35 days of age of broiler chickens (n=30 birds)

| Groups | G1 | G2 | G3 | G4 | G5 |
| --- | --- | --- | --- | --- | --- |
| Live wt.(g) | 2145.50<br>±1.20<sup>a</sup> | 2235.24<br>±1.70<sup>c</sup> | 2248.62<br>±1.80<sup>c</sup> | 2284.21<br>±1.10<sup>b</sup> | 2290.68<br>±1.85<sup>b</sup> |
| Carcass wt. (g) | 1523.31<br>±1.82<sup>a</sup> | 1612.80<br>±1.78<sup>c</sup> | 1630.51<br>±1.99<sup>c</sup> | 1650.71<br>±1.17<sup>b</sup> | 1667.03<br>±1.45<sup>b</sup> |
| Dressing (%) | 71.00 | 72.15 | 72.51 | 72.26 | 72.77 |
| Spleen (g) | 2.80±1.45<sup>d</sup> | 2.89±1.35<sup>c</sup> | 2.93±1.61<sup>b</sup> | 2.94±1.74<sup>a</sup> | 2.99±1.52<sup>a</sup> |
| Thymus (g) | 2.25±1.54<sup>d</sup> | 3.22±1.14<sup>d</sup> | 3.61±1.15<sup>c</sup> | 3.83 ±1.12<sup>b</sup> | 3.94±1.13<sup>a</sup> |
| Bursa (g) | 2.55±0.12<sup>d</sup> | 2.72±0.13<sup>c</sup> | 2.86±0.10<sup>b</sup> | 2.88±0.12<sup>b</sup> | 2.95±0.11<sup>a</sup> |
| Liver (g) | 44.38±1.83<sup>a</sup> | 44.50±1.71<sup>a</sup> | 45.35±1.95<sup>a</sup> | 45.15±1.35<sup>a</sup> | 45.28±1.45<sup>a</sup> |
| Heart (g) | 4.35±1.62<sup>a</sup> | 4.50 ±1.15<sup>a</sup> | 4.45±1.71<sup>a</sup> | 4.65±1.14<sup>a</sup> | 4.55±1.31<sup>a</sup> |
| Abdominal fat (g) | 2.0±0.02<sup>a</sup> | 1.80±0. 13<sup>b</sup> | 1.70±0.05<sup>b</sup> | 1.48±0.06<sup>c</sup> | 1.30±0.03<sup>d</sup> |

Means ± SD with different letter superscripts in each column are significant at P≤ 0.05
G1= Group of chicks fed on basal diet (BD)
G2= Group of chicks fed on BD supplemented with 500 g/ton of sodium butyrate (SB)
G3= Group of chicks fed on BD supplemented with 4 Kg/ton of rosemary leaves (RL)
G4 = Group of chicks fed on BD supplemented with 250 g/ton of SB + 2 kg/ton of RL
G5 = Group of chicks fed on BD supplemented with 500 g/ton of SB + 4 kg/ton of RL

## DISCUSSION

The current research shows that giving basal diets supplemented with SB increased BWG and FCR The findings **of Sikandar *et al*. (2017), Lan *et al*. (2020), Zhao *et al*. (2022), Kumar et al., (2023) and Deng *et al*. (2023)** were in agreement with our findings. The prior authors claimed that feeding basal diet supplemented with SB improved BWG and FER of broilers. On the contrary, dietary SB had no effect on the growth performance of broilers, according to **Zhang *et al*. (2011)**. The discrepancy between the two studies could be due to variations of added SB concentrations. While in this study a large quantity (500 g/ton) was added, the earlier authors used lower amounts (200 and 400 g/ton) of SB.

The results of this investigation showed that the rosemary herb improved the growth performance of broilers. Body weight gain (BWG) and feed conversion ratio (FCR) were both enhanced. The results of **Al-Kassie *et al*. (2008) and Yesilbag *et al*. (2011)** revealed that rosemary leaves increased growth performance in broilers. The findings of this study also agreed with those of **Mathlouthi *et al*. (2012); Loetscher *et al*. (2013); Rostami *et al*. (2015) and Ogwuegbu *et al*. (2021)**. Additionally, **Ghazalah and Ali, (2008)** reported a markedly higher growth rate and better feed conversion ratio when using rosemary powder at a 0.5% concentration in feed. According to **Petricevic *et al*. (2018)**, broiler diet supplemented with 0.4% dried and finely powdered rosemary leaves had a good impact on feed conversion and weight gain. The beneficial effects of utilizing various concentrations of rosemary powder in the feed were also confirmed by **Norouzi *et al*. (2015)**.

According to biochemical tests, adding SB and RL to broiler basal diets elevated serum levels of total protein (TP), albumin, globulin, superoxide dismutase (SOD), catalase (CAT), IgG and IgM but lowered levels of TC, TG, MDA, AST, and ALT. These results concurred with those of Lan *et al*. (2020) and Kumar *et al*. (2023) when using SB and with those of **Ghazalah and Ali, (2008) and Yao *et al*. (2023)** when using RL in feeds.

According to this study, adding SB and RL to the basal diet significantly raised macrophage phagocytic activity phagocytic index, lysozyme activity, and serum nitric oxide concentration in broiler chickens. The findings of **Sikandar *et al*. (2017); Lan *et al*. (2020); Ogwuegbu *et al*. (2021), and Sadurní *et al*. (2022)** were comparable to our findings. The previous authors came to the conclusion that supplementing basal diets with SB and RL at various concentrations enhanced immunoglobulin IgG and IgM and improved immunity in broiler chickens. Additionally, **Nafaa *et al*. (2023)** investigated how SB affected the intestinal immune response to Eimeria maxima infection and the histomorphological structure of the intestine of broiler chickens. The results showed that in the treated groups the intestinal villi and crypt depth were much longer than those of the control group. According to **Zhang *et al. (*2023) and Yao *et al*. (2023**), SB and RL exhibited considerably higher gut immunity than control birds against E. maxima infection. The authors concluded that adding RL to the basal diet raised the immunity of the ducks and broilers

The current findings reported that adding SB and RL to basal diets considerably improved carcass traits by increasing live weight, carcass weight, and dressing percent. Additionally, it reduced the weight of abdominal fat, while significantly increasing bursa, spleen and thymus weights. There were non-significant changes in the weight of heart and liver. These findings were in line with those of **Petricevic *et al*. (2018); Lan *et al*. (2020) and Ogwuegbu *et al*. (2021)** who discovered that adding SB and RL to the basal diet had a positive impact on carcass characteristics of broiler chickens.

## CONCLUSIONS

It is concluded that the best addition is to supplement the basal diet with 500 g/ton SB and 4 kg/ton R (group 5) for enhancing growth performance, normalizing biochemical indeses, boosting immune function, and producing good carcass traits of broiler chickens. Supplementation of basal diet with SB and RL produces growth-promotor, hypolipidemic, antioxidant, and immunostimulant effects. The mechanisms of action underlying these effects require further investigation in broiler chickens.

## DATA AVAILABILITY

The participants of this study do not give written consents for their data to be shared publicly, so due to the sensitive nature of the research supporting data are not available.

## AUTHOR CONTRIBUTIONS

Hamed Yahya Saifan conducted the experiments and collected data. Mostafa Abbas Shalaby, Khaled Abo-El-Sooud and Mohamd Ahmed Tony designed and supervised the work. Khaled Abo-El-Sooud performed statistical analysis. Mostafa Abbas Shalaby and Mohamed Ahmed Tony wrote original draft of the article and prepared the figures. Aya Mohye Yassin performed the biochemical analyses. Mostafa Abbas Shalaby wrote and revised the final manuscript before submission.

